# High-throughput, multiplexed quantification, and sorting of single EVs at single-molecule level

**DOI:** 10.1101/2024.10.31.621423

**Authors:** Juhwan Park, Michelle Feng, Jingbo Yang, Hanfei Shen, Zhiyuan Qin, Wei Guo, David A. Issadore

## Abstract

We have developed a platform for the high-throughput, multiplexed, and ultra-sensitive profiling of individual extracellular vesicles (EVs) directly in plasma, which we call BDEVS - Agarose **B**ead-based **D**igital Single Molecule-Single **EV S**orting. Unlike conventional approaches, BDEVS achieves single molecule sensitivity and moderate multiplexing (demonstrated 3-plex) without sacrificing the throughput (processing ten thousand of EVs per minute) necessary to resolve EVs directly in human plasma. Our platform integrates rolling circle amplification (RCA) of EV surface proteins, which are cleaved from single EVs, and amplified within agarose droplets, followed by flow cytometry-based readout and sorting, overcoming steric hindrance, non-specific binding, and the lack of quantitation of multiple proteins on EVs that have plagued earlier approaches. We evaluated the analytical capabilities of BDEVS through head-to-head comparison with gold-standard technologies, and demonstrated a ∼100x improvement in the limit of detection of EV subpopulations. We demonstrate the high throughput (∼100k beads / minute) profiling of individual EVs for key immune markers PD-L1, CD155, and the melanoma tumor marker TYRP-1, and showed that BDEVS can precisely quantify and sort EVs, offering unprecedented resolution for analyzing tumor-immune interactions and detecting rare EV subpopulations in complex clinical specimens. We demonstrate BDEVS’s potential as a transformative tool for EV-based diagnostics and therapeutic monitoring in the context of cancer immunology by analyzing plasma samples from patients with melanoma, where EV heterogeneity plays a critical role in disease progression and response to therapy.

**Graphical Abstract:** 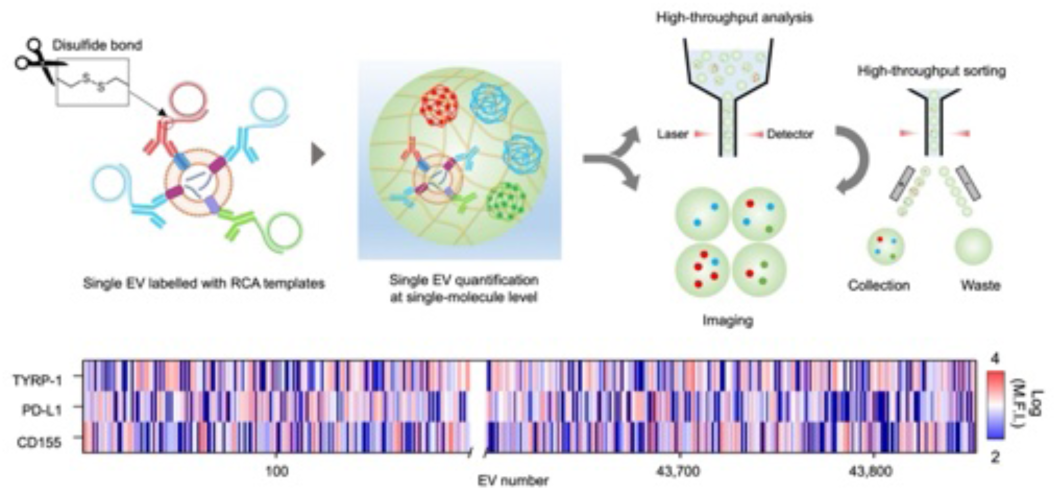

## Introduction

Extracellular vesicles (EVs), membrane-enclosed particles ranging from 30-1000 nm in diameter that are released by cells throughout our body and carry multiple molecular cargo from their mother cells into circulation, have attracted enormous attention for their diagnostic and therapeutic potential.[1–4] At the same time, there is a growing appreciation and understanding of the important mechanistic roles that EVs play in both health and disease, across a broad range of biology including cancer immunology,[5] infectious disease,[6] pregnancy,[7] and wound healing[8]. Unfortunately, limitations in the capability of current technology to resolve the heterogenous packaging of molecular cargo within individual EVs continues to hinder our ability to understand EV biology and to fully harness it for biomedical applications.[9,10] Conventional sample preparation and analyses that process EVs in bulk cannot resolve heterogeneity in EV composition and function; it has to be addressed at the single EV level. Technologies such as flow cytometry and single-cell sequencing, which can profile surface proteins on individual cells, have become essential tools in modern biology and have fueled progress in basic, as well as translational, science over the last several decades.[11] The development of analogous tools to profile individual EVs, however, poses a formidable, currently unsolved challenge. Because EVs have ∼100x smaller diameter than cells, and ∼*(100x)*^2^ less surface area, a technology that profiles EV surface markers generally requires 4 orders of magnitude greater sensitivity than that for cells. Indeed, recent studies have reported that EVs can have as few as single digit numbers of relevant surface proteins, requiring the quantification of EV surface marker expression down to the single molecule level.[12] Additionally, there are 10^11^ EVs present in each mL of blood, which is ∼10,000x than the number of cells in blood, requiring any technology for EVs to have a throughput that is ∼4 orders of magnitude greater than technology for cells. For example, for cancer diagnostics there are reportedly only 10^3^-10^4^ tumor derived EVs/mL of plasma, representing only 1 in 100 million EVs, thus requiring ∼10^9^ total EVs to be sampled in blood to observe a sufficient number of tumor derived EVs to not be limited by Poisson counting error.[13] Because sensitivity and throughput come at the expense of one another, building a system that can simultaneously achieve both the required sensitivity and throughput for EV analysis in clinical specimens, while also providing the level of multiplexing necessary to capture the relevant biology, has proven an obdurate challenge.

There has, however, been significant recent progress on the development of technology for single EV analysis, including platforms that have achieved single molecule quantification on individual EVs,[12,14–17] those that have sufficiently high throughput to quantify EVs in plasma[18–22], and those that sufficiently multiplex protein analysis on single EVs to capture relevant biology. [23–26]. Many of the recently developed techniques to resolve single EVs are based on fluorescence imaging, which has been successful in resolving individual EVs and performing multiplexed analysis of their cargo, including both protein and RNA.[27–31] However, these imaging modalities are limited to analyzing the EVs present in a field of view, typically <10^4^, which is several orders of magnitude less than what is required to resolve rare EVs in blood. Notable amongst the recent work in this area, a cyclic fluorescence imaging of captured EVs was recently reported that achieved the multiplexed analysis of up to 12 biomarkers ∼10,000 individual EVs.[26] Moreover, conventional readout strategies of fluorescence immunolabeling, such as imaging or flow cytometry lack the sensitivity to resolve single digit quantities of fluorescent labels present on EVs, and chemical amplification of the signal prior to measurement can aid high throughput readout with single molecule sensitivity. However, many amplification strategies, such as rolling circle amplification (RCA)[15] or hybridization chain reaction (HCR)[21] that grow a nucleic acid structure on the immunolabel are limited by steric hindrance on EVs, because these products are ∼10x larger than the EVs themselves. Droplet microfluidics, where single EVs are loaded into droplets provide a solution to this problem by providing a container to retain the free-floating amplified products released from an EV. In droplet-based approaches, surface markers on EVs are either barcoded for downstream sequencing analysis or labeled with an enzyme for digital enzyme-linked immunosorbent assay (dELISA). While recent dELISA platforms have been successful in resolving rare EVs in plasma, LOD < 10 EVs/ µL,[18,19] they are limited in that they can only count the number of droplets positive for a particular pair of biomarkers and cannot quantitate their expression level. Droplet-based DNA barcoded sequencing approaches can be highly multiplexed, but have only been used at relatively low throughput ∼10^4^ EVs. In one particularly noteworthy effort, as many as 38 biomarkers on ∼13,000 individual EVs were quantified.[25] Despite this recent progress in the field, a single platform has yet to be developed that can combine the sensitivity to detect single molecules on individual EVs, the multiplexing to resolve relevant EV subsets based on surface protein expression, and the throughput to make it relevant to the study of EV subpopulations in complex clinical specimens such as blood.

To address this challenge, we present a new technology tailored specifically to resolve rare EV populations directly in plasma that can profile individual EVs at high-throughput (∼10,000 of EVs/min), with moderate multiplexing (3-5plex), and achieve ultra-sensitive digital protein detection on each EV. We coin this technique Agarose **B**ead based **D**igital Single Molecule - Single **EV S**orting (BDEVS). To achieve this performance our platform combines several innovations: 1. BDEVS achieve single molecule sensitivity without sacrificing throughput by amplifying each targeted protein on the EV using RCA.[15,32–35] We avoid the issue of steric hindrance that can plague signal amplification of surface proteins by *in situ* cleavage of the RCA templates from the EV, such that the RCA products are dispersed over a microscale sized bead allowing digital counting of RCA products. 3. We can process and readout BDEVS at high throughput using conventional laboratory equipment (i.e. centrifuges, flow cytometer), by loading each EV into a microscale solid agarose bead and labelling the retained RCA products associated with individual surface proteins, which would be prohibitively cumbersome to accomplish using conventional water-in-oil or double emulsion droplet microfluidics.[36,37] In a related recent work from our lab, we have used agarose microbeads to analyze individual mitochondrial DNA (mtDNA) within individual cells by lysing the trapped cells inside agarose microbeads, followed by padlock probe-based RCA of the mtDNA. This prior approach trapped the mtDNA (16kb) within each microbead based on its large size relative to the agar microgels pores.[32] In contrast, the RCA templates associated with biomarkers of single EV used in BDEVS (<100 bp) are too small to be retained by our agarose beads. To address this challenge, we perform RCA directly in the emulsion phase, generating RCA products sufficiently large to be retained within the microgel for downstream processing. Additionally, unlike many alternative approaches,[18,19,24] BDEVS does not use antibody functionalized microbeads, which greatly reduces its background signal from non-specific binding of Abs, as the surface area of a 5 µm bead is ∼25,000x larger than that of an EV, proportionally reducing the probability of non-specific labeling.

While BDEVS is broadly applicable across many applications (e.g. neurodegenerative diseases, diabetes), in this work we focus on tumor-immune interaction in melanoma as a testing ground for our technology. We chose tumor immunology to develop and test our technology because EV heterogeneity is already known to play a key role in disease progression and therapy response,[5,38–40] is relatively better studied in cancer than in other fields, and there are numerous well characterized protein markers, and practically, antibodies for these markers. In this study, we use BDEVS to profile single EVs for the immune markers PD-L1 and CD155 and the melanoma tissue specific marker TYRP-1 with high-throughput and multiplexed quantification from a plasma sample. To this end, we first quantified the analytical performance of BDEVS by quantifying multiple tetraspanins (CD9, CD63, CD81) on individual *in vitro* derived EVs and performed a a head-to-head comparison with a low throughput gold standard technology (EXOVIEW). We additionally demonstrated that in addition to profiling the microbeads at high throughput using flow cytometry, we could also sort single EVs by using FACS and defining a “gate” based on our multiplex RCA signal, and analyze the sorted beads downstream with fluorescence microscopy for single molecule analysis on each individual EV by counting individual fluorescently labeled RCA products. Subsequently, we validated our BDEVS assay for cancer immunology markers PD-L1 and CD155, and TYRP-1, using both *in vitro* systems as well as a set of plasma samples from a cohort of patients with melanoma (n=15). We validated that we could profile EVs with both single EV and single molecule resolution even in samples with EV signals below the noise floor for conventional ELISA.

## Result

The BDEVS workflow is designed to be robust and easily carried out using commonly available laboratory equipment, besides the microfluidic chip used to generate the agarose droplets, and analyzed at high throughput using either flow cytometry, imaging for single molecule quantification, or flow sorting to allow downstream processing of selected EV subpopulations (**Fig. 1**). In our workflow, first, the EVs in a sample are incubated with primer-conjugated antibodies for 1 h at room temperature. To ensure a constant number of primer per antibody, we incorporated oYo link technologies (AlphaThera) that conjugate DNA oligomer to the Fc region of antibody after UV curing.[41] By incorporating oYo link technologies, we are able to quantify the number of biomarker expressed on single EV based on the number of RCA products per agarose bead (**Table S1**). After labelling EVs with detection antibody-DNA primer, unbound antibody conjugates are washed away using a size exclusion chromatography (SEC) column (IZON). These EVs are subsequently labelled with 50 nM circle DNA (pre-circularized by ssDNA circLigase kit, Lucigen) for 2 h at room temperature. Using a parallelized flow focusing droplet generator operated above the melting temperature of agarose (**Figure S1**),[42] these EVs are encapsulated into 0.33% agarose droplets with 3x RCA mixture in a continuous phase of HFE-7500 (10 mL/h), at high throughput (>20,000 drops per min). The agarose droplets are collected in an ice bucket for immediate gelation, and subsequently incubated at 37 °C for 2 h for the RCA reaction. During this step, the disulfide bond that we incorporate between the oYo link and DNA primer is cleaved by a reducing agent that we include in our RCA mix, releasing the RCA product from the EV and avoiding steric hindrance. We then transfer the agarose beads to an aqueous phase (See Methods). The RCA products are labelled with fluorescence detection probes. After washing away unbound detection probes three times with PBST by centrifugation, agarose beads are analyzed by imaging, flow cytometry, or FACS.

**Figure 1.**
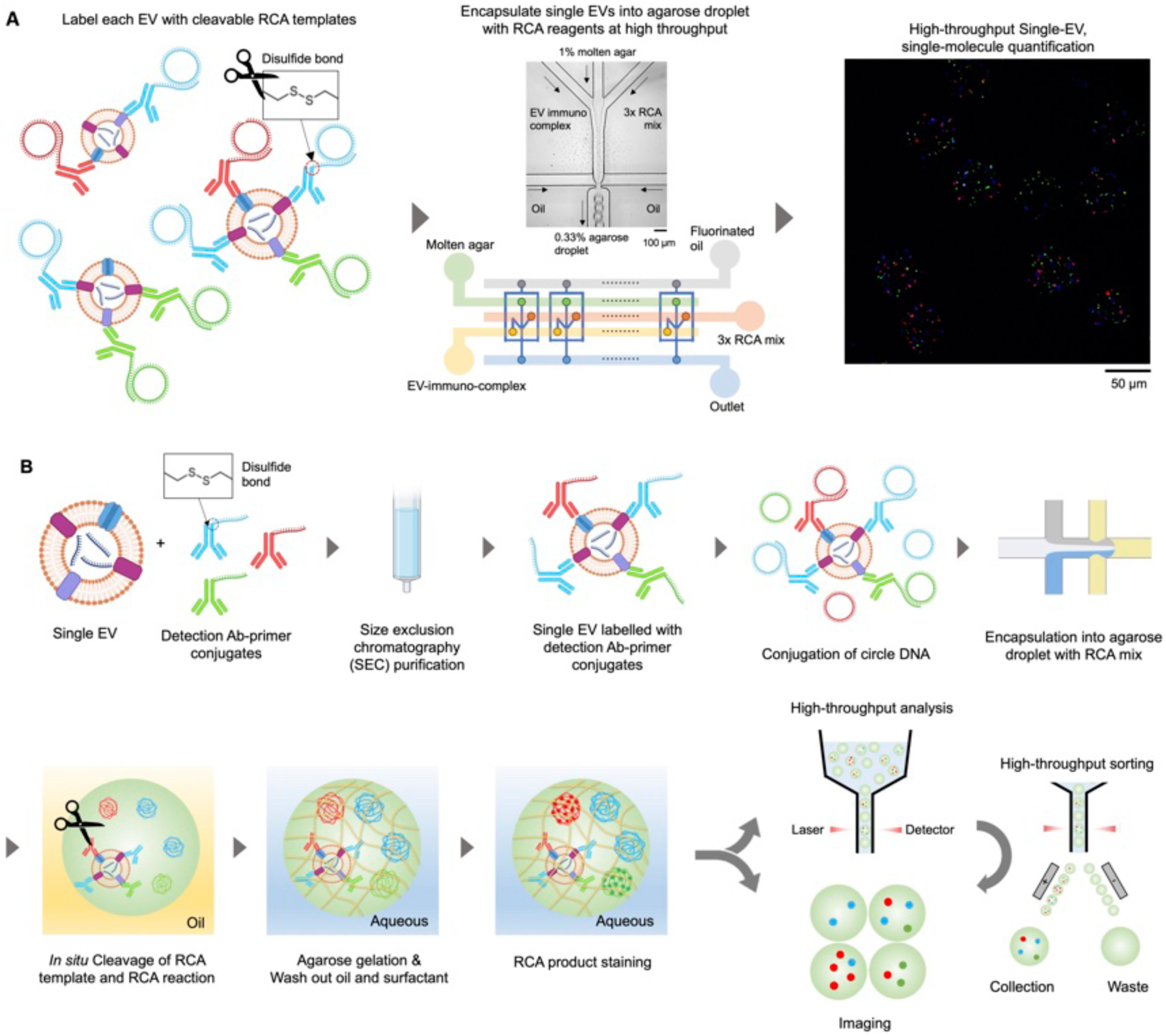
Workflow for BDEVS. (A) Single EVs labelled with antibody-RCA template are encapsulated into agarose droplets along with RCA mixture. RCA products are visualized within agarose beads to quantify multiple biomarkers on single EVs. (B) Experimental procedures for BDEVS. Once encapsulation of single EVs labelled with RCA templates are encapsulated into agarose droplets along with RCA mix, the RCA templates are cleaved from the EV and the RCA reaction is performed simultaneously. After the RCA reaction, agarose droplets are gelated and transferred into the aqueous phase followed by labelling of RCA products with detection probes. After the assay, single EVs are sorted by flow cytometry and analyzed using post-sort fluorescence imaging.

We first characterized the BDEVS assay using pre-formed RCA templates that consist of 30 bp primer hybridized to 68 bp circle DNA (**Fig 2A**). The elevated temperature used to melt the agarose for microfluidic droplet formation must be above the gelling point of agar (T > 20 °C) but not so high that it denatures the enzyme (phi29 DNA polymerase) (T < 65 °C) used in RCA. To optimize the operating temperature, we performed our BDEVS assay at droplet generation temperatures of 25, 30, 40, and 50 °C. At all temperatures we were able to generate monodisperse agarose droplets (CV < 6%) (**Figure S2**), but we found that the RCA efficiency decreases as temperature increases over 40 °C (**Fig. 2B**). Agarose beads with RCA products were analyzed using flow cytometry (BD Science A3 Lite, Penn flow core) (**Figure S3**). Flow cytometry confirmed RCA efficiency is hindered by temperatures > 40 °C (**Fig. 2C**). Based on these experiments, we performed droplet generation at 30 °C. To quantitatively evaluate the performance of BDEVS, we next tested it using various known concentration of RCA templates. Fluorescence microscope imaging (**Fig. 2D**) and flow cytometry (**Fig. 2E**) confirmed that the mean fluorescence intensity (MFI) of the beads was proportional to the concentration of the RCA templates from 500 fM to 50 pM concentration (**Fig. 2E**). To increase the sensitivity of the assay, such that even one single RCA product would be detected, the diameter of agarose droplet was decreased to 27 µm (CV = 8%) from 61 µm (CV = 5%) to improve the signal-to-noise ratio (**Figure S4**). Using these 27 µm diameter droplets, at 50 fM of RCA template, we confirmed that the RCA products were digitally distributed over agarose beads by noting less than 10% of agarose beads are detected as a positive (**Fig. 2F -inset**). Using this same concentration of RCA template, we confirmed that a droplet with a single template can be differentiated from a bead with zero templates (**Fig. 2F**). We next titrated the RCA template concentration and measured the fraction of beads that were positively fluorescent in flow cytometry to determine the analytical performance of BDEVS in detecting RCA templates (**Fig. 2G**). Limit of detection (LOD) and limit of quantification (LOQ) were found to be 443 aM and 846 aM, respectively, and a dynamic range (DR) of 1196. We note that the DR can be expanded by increasing the number of beads analyzed or by expanding beyond the digital regime by quantifying the fluorescence intensity of the positive beads. Key to achieving this performance was the analysis of > 0.5 million beads per measurement, such that the variance in the measurement of the blank signal was not dominated by Poisson noise. The blank signal had a false positive rate of only 0.005 %, and at 0.5 million beads we measured approximately 25 beads, resulting in a Poisson noise √“/” < 20%. We characterized BDEVS’s capability for multiplexed detection of RCA product. We loaded droplets with three different RCA templates, each with an associated complementary detection probe, labeled with a different fluorophore (Alexa Fluor 488, ATTO565, ATTO647). In this experiment, 5 pM of each RCA template was encapsulated into the agarose droplets for the BDEVS assay, and under inspection by fluorescence microscopy distinct RCA products can be resolved for each of the RCA templates (**Fig. 2H**). We compared multiplex BDEVS with singleplex assay, keeping the concentration of each of the RCA templates constant, to evaluate whether there was any hindrance or crosstalk in the RCA reactions (**Fig. 2I-K**). MFI measured in the multiplex assay changed only marginally compared to the respective singleplex assays, with shifts in MFI less than the CV of the singleplex assay, demonstrating minimum cross-talk in the multiplex assay.

**Figure 2.**
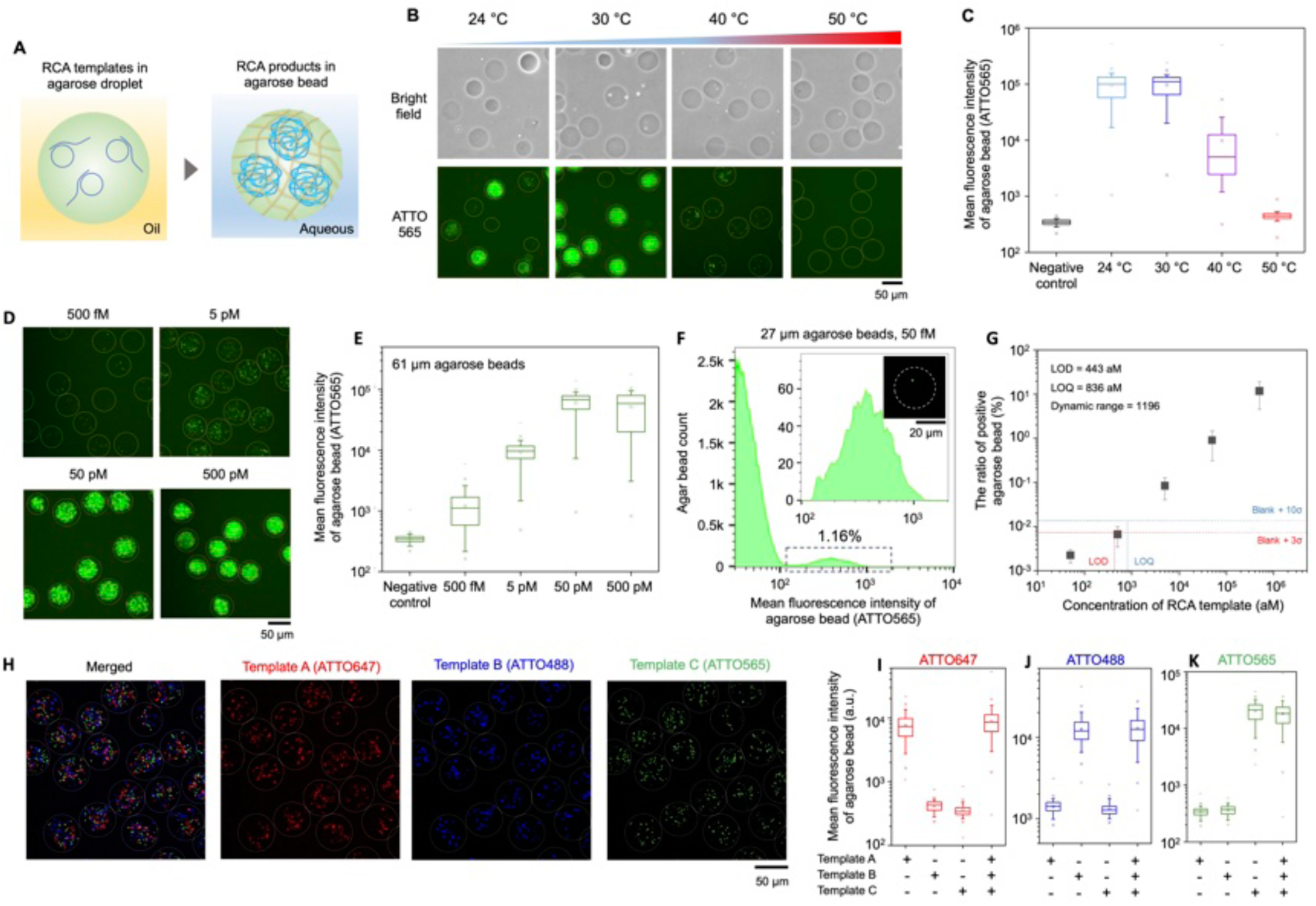
Characterization of agarose droplet-based RCA. (A) Pre-formed RCA templates were encapsulated into agarose droplet with the RCA mix. After the RCA reaction, agarose gelation, and transferring to aqueous phase, RCA products within agarose beads were labelled with fluorescence detection probes. (B) Temperature for agarose droplet generation was optimized to avoid phi29 polymerase inhibition while generating monodisperse agarose droplets. Micrographs shows the RCA products within agarose beads generated at various temperatures. (C) Mean fluorescence intensity (MFI) of agarose beads generated at different temperatures were analyzed using flow cytometry and compared with a negative control without RCA templates. (D) Fluorescence micrographs of agarose microbeads demonstrate the the RCA product count changes proportionally to the RCA template concentration. (E) MFI of agarose beads with different concentration of RCA templates were analyzed using flow cytometry. MFI saturation occurred around at 50 pM due to the limited agarose droplet volume. (F) Comparison of analysis of RCA products using flow cytometer and fluorescence imaging demonstrates that a single RCA product in an agarose bead could be detected. (G) Positive rate of agarose beads was assessed with respect to the concentration of RCA template to validate the analytical performance of the agarose droplet based RCA assay. (H) A mixture of three different RCA templates was encapsulated into each agarose droplet to validate multiplexed RCA. Fluorescence micrographs show three different RCA products are distributed within agarose beads. (I-K) Multiplex RCA for three different RCA templates was compared to singleplex RCA for each template, indicating the independence of each RCA reaction.

We next evaluated BDEVS’s capability to measure single EVs using Mel-624-B7H1 (generated by Dong, et al.,[43] by transfecting a mel-624 with a B7-H1 expression (Mayo Clinic)) cell culture derived EVs. To evalulate the analytical performance of BDEVS, we first evaluated a singleplex BDEVS assay for the tetraspanin CD63 (**Fig. 3A**). We chose CD63 because it was found to be the most abundant tetraspanin marker based on EXOVIEW measurements (**Figure S5**). We first titrated the concentration of EVs, measured using nanoparticle tracking analysis (NTA)(Zetaview), to find the concentration at which the EVs are digitally loaded into the agarose droplets (<0.1 EV/bead) (**Fig. 3B**). We evaluated the spatial distribution of the RCA products within the agarose bead, to confirm that the RCA products are released from the surface of the EV using our cleavable linker (**Fig. 3C**). The distribution of the RCA products was indeed uniformly distributed across the bead, whereas in a negative control when the reducing agent is not included in the RCA mix, RCA products are concentrated at the location of the EV, confirming the utility of our cleaving to distribute the RCA products within the agar microbeads so that they can be resolved for accurate quantitation. We next evaluated the capability to quantify the BDEVS at high throughput using flow cytometry (**Fig. 3D**). We quantified the fraction of positive beads with any RCA products, i.e. the average number of EVs per bead (AEVB), and first confirmed that we are operating in the digital regime (AEVB < 10%). To validate that this signal was indeed from single EVs, we performed various negative control experiments (**Fig.3E**). In separate experiments, we sequentially stripped away key elements of the BDEVS assay. First, using a high concentration input of the Mel-624-b7h1 cell line derived EVs (1,500 EVs/µL), we replaced the CD63 antibody with an isotype control (Mouse IgG1, κ), we removed the Ab-primer, using only the circle DNA, and we removed the circle template as well. Finally, we used the full BDEVS assay but spiked zero EVs from the Mel-624-B7H1 cell line. For all negative controls, we measured a negative result, and measured AEVB for isotype, no EV, circle DNA, and PBS control to be 0.13%, 0.035%, 0.05%, and 0.009%, respectively. Based on these results, the background signal arises predominantly from non-specific binding of the antibody-primer to EVs. We next titrated the number of Mel-624-B7H1-derived EVs and quantified the AEVB to determine the analytical performance of BDEVS in detecting single EVs. A limitation of this study is that our ground truth concentration was defined using NTA, which reports the total number of particles that are appropriately sized, whereas BDEVS reports only those EVs that are positive for CD63. As such, the reported performance metrics are somewhat underestimated. However, EXOVIEW estimates that > 60 % of EVs are positive for CD63, and so our underestimation of our performance is small (**Figure S5**). EVs labelled with RCA templates purified by SEC were titrated to assess performance of the assay. The threshold in flow cytometry was set to select beads that contain more than one RCA product, to best distinguish true EVs from background (Threshold = 1,000). LOD and LOQ were found to be 7.0 EVs/µL and 19 EVs/µL, and DR = 790 after the optimization of the assay condition, similar to the best performances reported by digital ELISA based platforms (**Fig. 3F, Figure S6**).[18] We note that this performance comes from the combination of the low false positivity rate of our blank (AEVB_b_= 0.0003%) and measuring enough beads (>500,000 per assay) such that the performance is not Poisson noise limited. Using a high input of EVs, 1,500 EVs/μL, we compared the measured AEVB = 0.196%, with both the no EV control (AEVB = 0.0012%) and an isotype control (AEVB = 0.0028%) (**Fig. 3G**). When the threshold for distinguishing EVs with CD63 to detect more than one RCA product was applied, the AEVB from no EV and the isotype control decreased by 29 and 46 times, respectively, compared to when the threshold was set at 100 (**Fig. 3E**). Compared to the conventional method that captures EVs using microbeads followed by immune-RCA labelling, BDEVS showed >1,000 times lower background signal when there were no target EVs (**Figure S7**).

**Figure 3.**
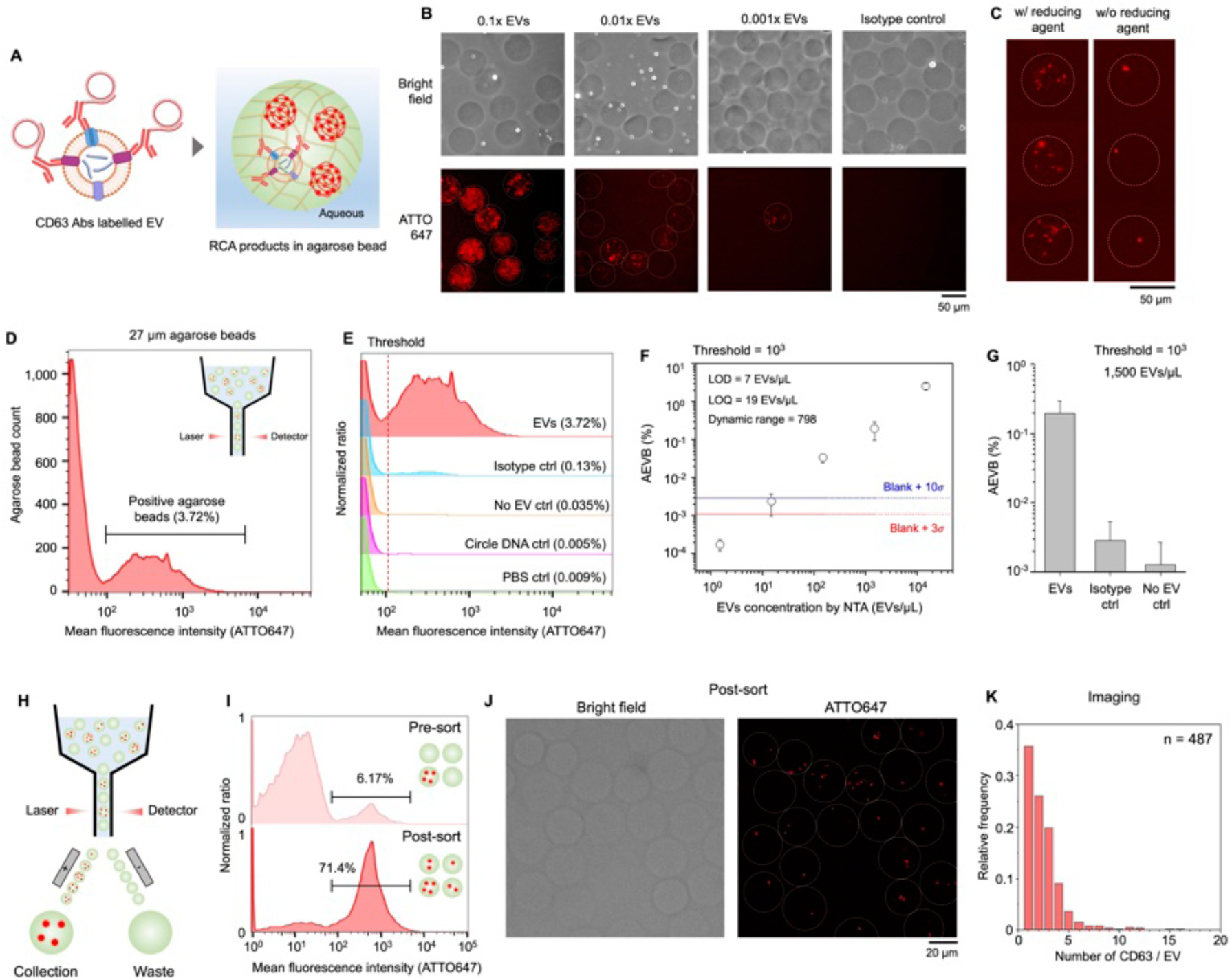
Quantitative single-molecule assay of single EVs. (A) of workflow to analyze single EVs for CD63 expression using BDEVS. (B) Micrographs of beads in which we have performed the BDEVS assay on single EVs. As dilution factor increases, we reach the digital regime, where RCA products generated from a single EV are distributed throughout the agarose bead. (C) The introduction of reducing agent overcomes the steric hindrance of multiple RCA reactions on the surface of a single EV, enabling recognition of single surface markers. (D) MFI of agarose beads were analyzed using flow cytometry. The measured average number of EVs per bead (AEVB) was below 10%, verifying operation in the digital regime. (E) AEVB measured for isotype, no EVs, circle DNA only, and PBS control experiments. (F) Titration curve for EVs labelled with antibody-RCA templates. (G) AEVB compared with isotype and no EVs control at the same concentration of EVs and the same dilution factor. (H) Agarose beads with RCA products were sorted using flow cytometry to improve the throughput of fluorescence imaging. (I) AEVB of agarose beads before and after the sorting, showing that the positive agarose beads are concentrated > 10x. (J) Micrographs showing agarose beads after the sorting. (K) A histogram showing the distribution for the number of CD63 molecule per EV.

While flow cytometry allows for high throughput analysis of EVs (>1000 beads/sec), it cannot perform the precise counting of RCA products that we achieve in low throughput microscopy. To bridge this gap, we demonstrate that it is possible to perform fluorescence activated sorting of the agarose beads for downstream imaging. We demonstrated compatibility with conventional laboratory equipment by sorting using a Symphony-S6 FACS from Penn flow core (BD bioscience) (**Fig. 3H,I**), sorting based on positive signal for CD63, and analyzing the sorted beads downstream using fluorescence microscopy to count individual CD63 associated RCA products for each EV. We observed on average 2.5 RCA products associated with CD63 per bead (CV = 89.9%) (**Fig. 3J,K**). In this experiment, 25,919 CD63 positive EVs within agarose beads were sorted from 442,993 agarose beads (42 beads per second), and 487 agarose beads were imaged using a low magnification (20x) microscope.

We next evaluated BDEVS’s capability to perform multiplex quantification of the tetraspanins CD9, CD63, CD81 on EVs derived from Mel-624-b7h1 cell culture media (**Fig. 4A**). First, using a flow cytometry we validated a digital distribution of EVs in the agarose beads (AEVB < 10%), where an EV is defined as being positive for either CD9, CD63, CD81, or any combination of the three (**Fig. 4B**). We quantified CD9, CD63, and CD81 expression of individual EVs at high throughput, analyzing in total > 60,000 EVs with > 8,800 EVs / minute throughput (**Fig. 4C**). The ratio of EVs co-localized with CD63, CD81, CD9, CD63/CD81, CD9/CD63, CD9/CD81, CD63/CD81/CD9 were 61.1%, 19%, 5%, 13.1%, 1%, 0.2%, and 0.6%, respectively (**Fig. 4D**). Co-localization of tetraspanin and the number of tetraspanins per single EV were further analyzed by imaging after sorting agarose beads having any RCA products (**Fig. 4E,F**). Agarose beads having any RCA products associated with CD63, CD81, or CD9 were sorted using FACS (Symphony-S6). Using these sorted beads, RCA products for the multiplexed assay can be quantified at the single molecule level using fluorescence microscopy (**Fig. 4G**). We found that on average 2.7 tetraspanin molecules (CV = 100%) were expressed per single EV with high heterogeneity. Both imaging and flow cytometry showed heterogeneity of tetraspanin expression on EVs, and the results of them qualitatively agreed each other (**Figure S8**). We compared the heterogeneity of tetraspanin expression that we measured with BDEVS to the commercial EXOVIEW system and found qualitative agreement (**Fig. 4H,I**). Both results agreed that CD63 is the most abundant tetraspanin, and CD9 is the least expressed tetraspanin. Additionally, the MFI of positive agarose beads for each fluorescence channel was compared between single-plex assays of each target molecule and the multiplex assay to validate if there was steric hindrance due to the molecular crowding (**Fig. 4J-L**). There was no significant difference in MFI between single-plex and multiplex assays, confirming that the labelling of EVs was not hindered by multiplexing of the assay.

**Figure 4.**
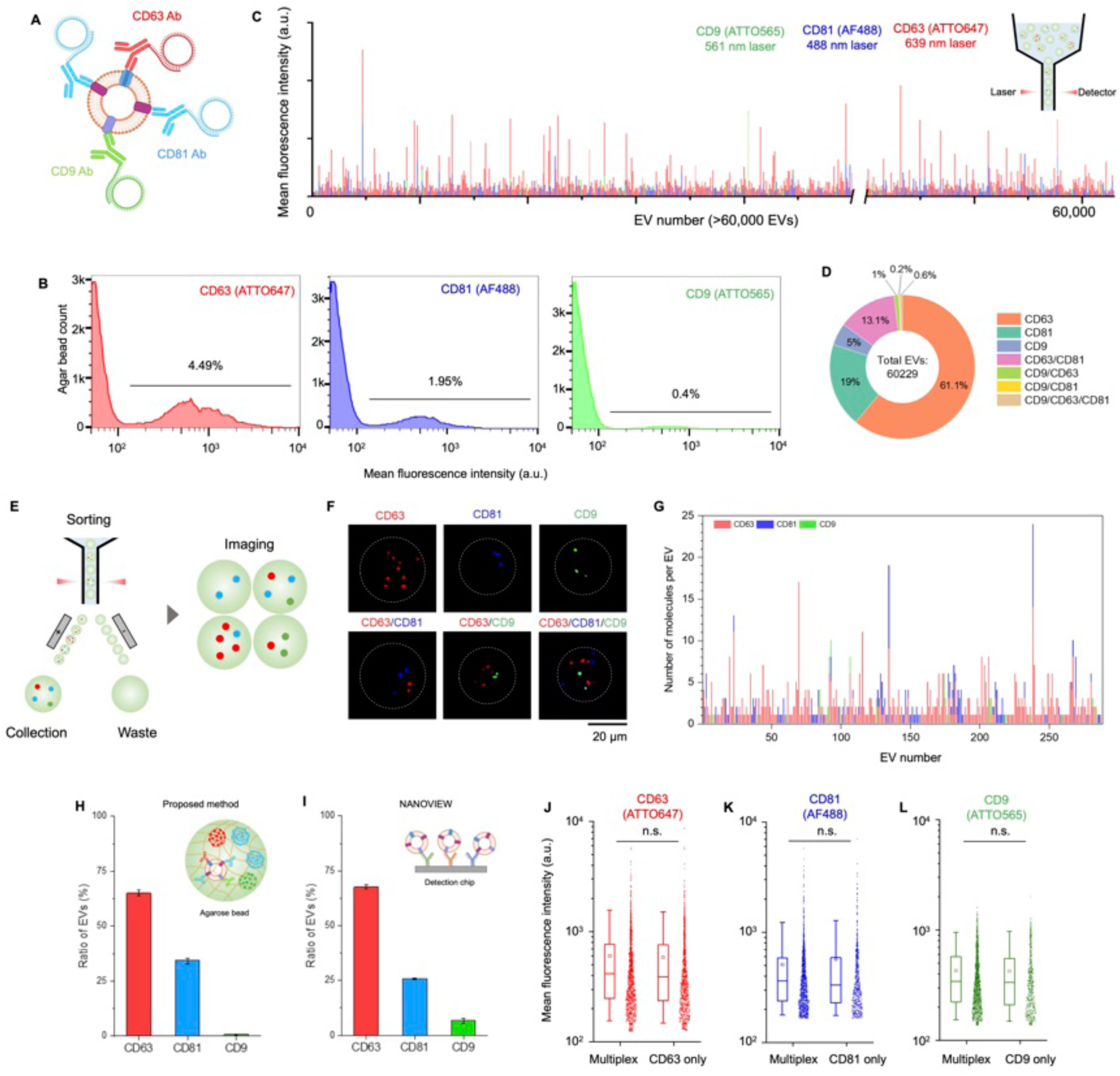
Quantitative and multiplexed single EV, single-molecule assay for tetraspanin analysis. (A) A schematic of workflow to analyze single EVs for CD9, CD63, and CD81 expression using BDEVS. (B) A histograms of AEVB in each fluorescence channel showed that AEVB was below 10% for each channel, verifying operation in the digital regime. (C) MFI of three different single-EV RCA products for > 60,000 EVs analyzed using flow cytometry. (D) Ring plot of the proportion of various population of EVs based on co-localization of tetraspanin. (E) RCA products labelled with each fluorescence detection probe (Alexa fluor 488, ATTO565, ATTO647) within a single agarose bead were counted after flow sorting. (F) Micrographs taken after the flow sorting shows various population of EVs co-localized with different tetraspanin. (G) The number of tetraspnin per EV was analyzed by fluorescence imaging after the flow sorting. Ratio of EVs detected by each tetraspanin antibody was compared between (H) BDEVS and (I) EXOVIEW, which shows CD63 is the most abundant tetraspanin and CD9 is the least expressed. (J-L) MFI of agarose beads with single EV was compared between multiplex and singleplex assay for each tetraspanin, which indicates there is little steric hindrance in multiplex labelling by noting no significant signal decrease in the multiplex assay.

Moving beyond tetraspanins, we next evaluated whether BDEVS can be applied to measure EV biomarkers relevant to cancer immunology. We measured TYRP-1, a melanoma specific biomarker, as well as the immune checkpoint proteins PD-L1 and CD155, immune markers at the single EV level (**Fig. 5A**). There has been a particular interest in studying PD-L1, a protein expressed on both cancer cells and their EVs [38,44], and its role in helping tumors evade the body’s immune response in cancers such as melanoma.[43] In healthy tissue, the expression of PD-L1 molecules on cell surfaces is used to protect cells from an immune attack.[45,46] In cancer, this process is exploited by tumor cells that overexpress PD-L1, both on the surface of cells and also on EVs released to the tumor microenvironment and circulation[38,47], where they interact with PD-1 on the surface of cytotoxic T cells and lead to their exhaustion and apoptosis[46]. Because of this role in cancer immune interaction, PD-L1 on EVs has been heavily studied as a predictor of the outcomes of immune check point blockade-based therapies in cancers such as melanoma, gastric, and lung cancers.[38,48–50] In addition to PD-L1, other immune markers such as CD155 (a.k.a. PVR) have also been shown to be associated with anti-PD1 Immunotherapy in metastatic melanoma, and may be beneficial biomarkers to predict personalized treatment response.[51–54] Previous efforts to profile the interaction between tumor cells and the immune system through EVs have been hindered by technological limitations, such as the inability to analyze individual EVs, insufficient throughput for detecting rare PD-L1+ EVs in blood, and the inability to quantify protein expression, multiplex immune markers, or determine EV origin.[19] We first evaluated BDEVS on EVs isolated from mel-624-B7H1 cell culture medium. The AEVB measured for TYRP-1, CD155, and PD-L1 was 2.62%, 3.04%, 3.19%, respectively (**Fig. 5B**). Co-localization of biomarkers was analyzed using flow cytometry (**Fig. 5C**), and we found co expression of TYRP-1/CD155, TYRP-1/PD-L1, CD155/PD-L1, and TYRP-1/CD155/PD-L1 was 2%, 1.4%, 1.8, 0.6%, respectively, relative to the total number of EVs measured (**Fig. 5D**). Agarose beads that had at least one RCA product were sorted using FACS for imaging. Based on imaging analysis, average 2.1 PD-L1 (CV = 58%), 1 TYRP-1 (CV = 56%), 0.8 CD155 (CV = 54%) was expressed per single EV (**Table S1**).

**Figure 5.**
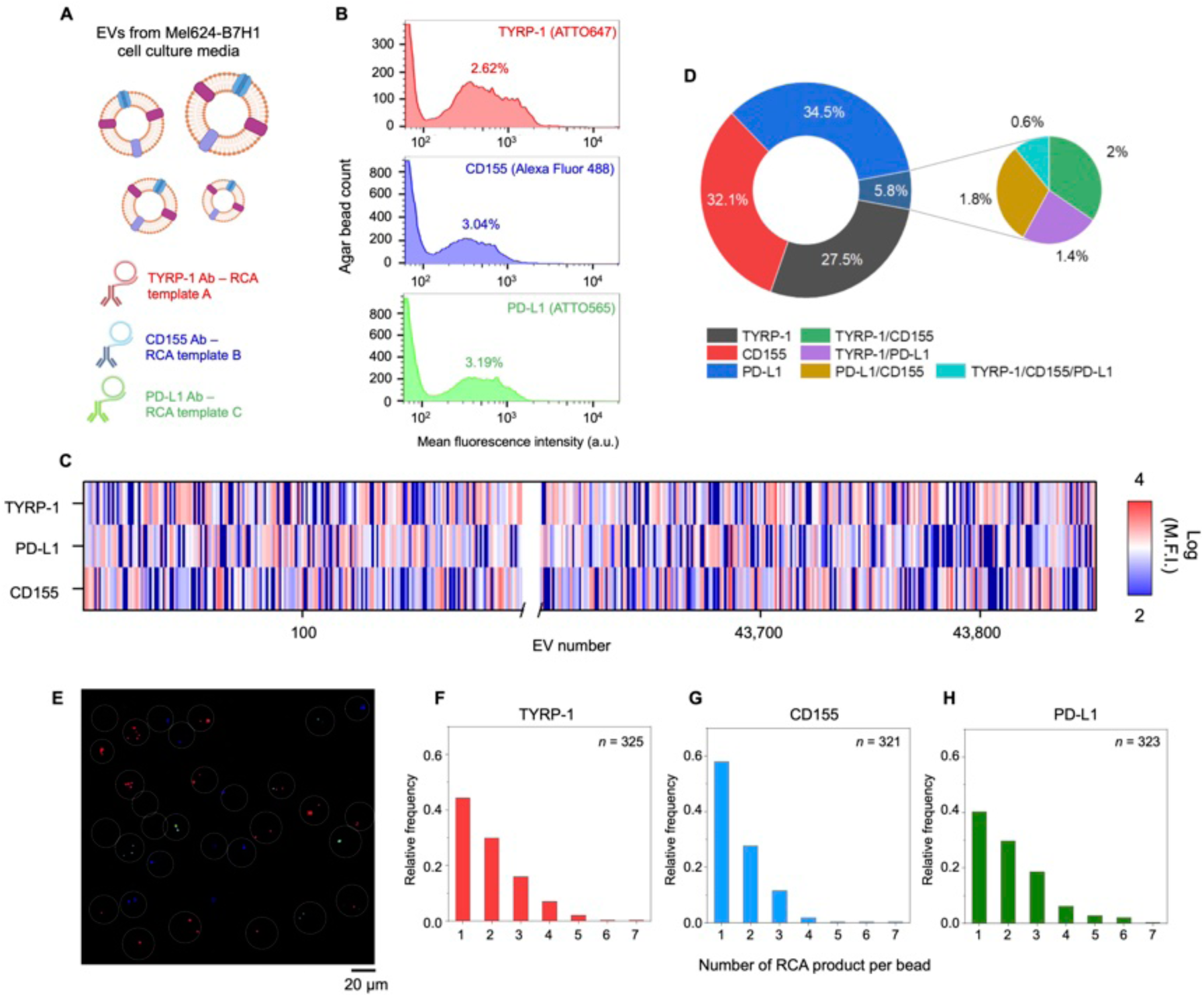
Analysis of biomarkers relevant to cancer immunology in single EVs. (A) Schematic of workflow to analyze single EVs from Mel-624-B7H1 cell culture for TYRP-1, CD155, and PD-L1 expression using BDEVS. (B) Histograms of the results of flow cytometry analysis quantifying the positive rate of agarose beads for each biomarker. (C) Heat map showing exosomal TYRP-1, CD155, and PD-L1 expression level that isolated from Mel-624-B7H1 cell culture medium. (D) Ring plot showing the proportion of different EVs populations. < 6% of EVs are co-localized with more than 1 biomarker. (E) Micrographs of agarose beads with RCA products after the flow cytometry sorting. (F-H). Histograms showing the number of RCA product per bead that represent expressed number of each biomarker per EV.

Subsequently, we tested our BDEVS assay for TYRP-1, PD-L1, and CD155 in single EVs in a cohort of N = 15 melanoma patients’ plasma samples (volume = 15 µL). Single EVs from 10x diluted plasma sample were labelled with each detection antibody conjugated with primer and were purified by SEC and labelled with circular DNA for the agarose droplet encapsulation with RCA mix (See method, **Fig. 6A**). To validate BDEVS can detect rare population of EVs relevant to cancer immunology from plasma samples with many background EVs, first pooled plasma from 5 melanoma patients was analyzed to compare the positive rate of agarose beads between target and isotype antibodies. Each target antibody showed 131 (TYRP-1), 31 (CD155), 75 (PD-L1) times higher positive rate than isotype control in BDEVS whereas ELISA showed 0.69 (TYRP-1 / PD-L1), 1.08 (CD155 / PD-L1) times higher signal intensity with target antibodies in comparison with isotype control (**Figure S9**). Then we analyzed EVs from each plasma sample of 15 melanoma patients (**Fig. 6B**). The ability of BDEVS to detect rare population of cancer immunology relevant EVs from plasma was compared to sandwich ELISA (**Fig. 6C-F**). The ratio of CD155/PD-L1 or TYRP-1/PD-L1 dual positive agarose bead was measured by BDEVS, and the fluorescence signal from CD155/PD-L1 or TYRP-1/PD-L1 dual positive EVs was measured by ELISA. BDEVS was able to detect positive EVs, above the LOD of our system, in 100% of the patient samples, reporting the fraction of beads that are dual positive for CD155/PD-L1 and TYRP-1/PD-L1, allowing direct comparison to respective conventional sandwich ELISAs. Conventional sandwich ELISA for CD155/PD-L1 and TYRP-1/PD-L1 were only able to detect signals above LOD for 33% and 6.7% (**Fig. 6G**). To demonstrate BDEV’s capability not only to count biomarker-specific EVs but also to quantify biomarker expression levels on individual biomarker-positive EVs, the mean fluorescence intensity of agarose bead populations, gated in flow cytometry analysis, was compared for each biomarker (PD-L1, CD155, and TYRP-1) (**Fig. 6H-J**). Furthermore, agarose beads were sorted and imaged to quantify the number of biomolecules per extracellular vesicle (EV) from samples of 5 melanoma patients, revealing variability in biomarker levels between individuals (**Figure S10**).

**Figure 6.**
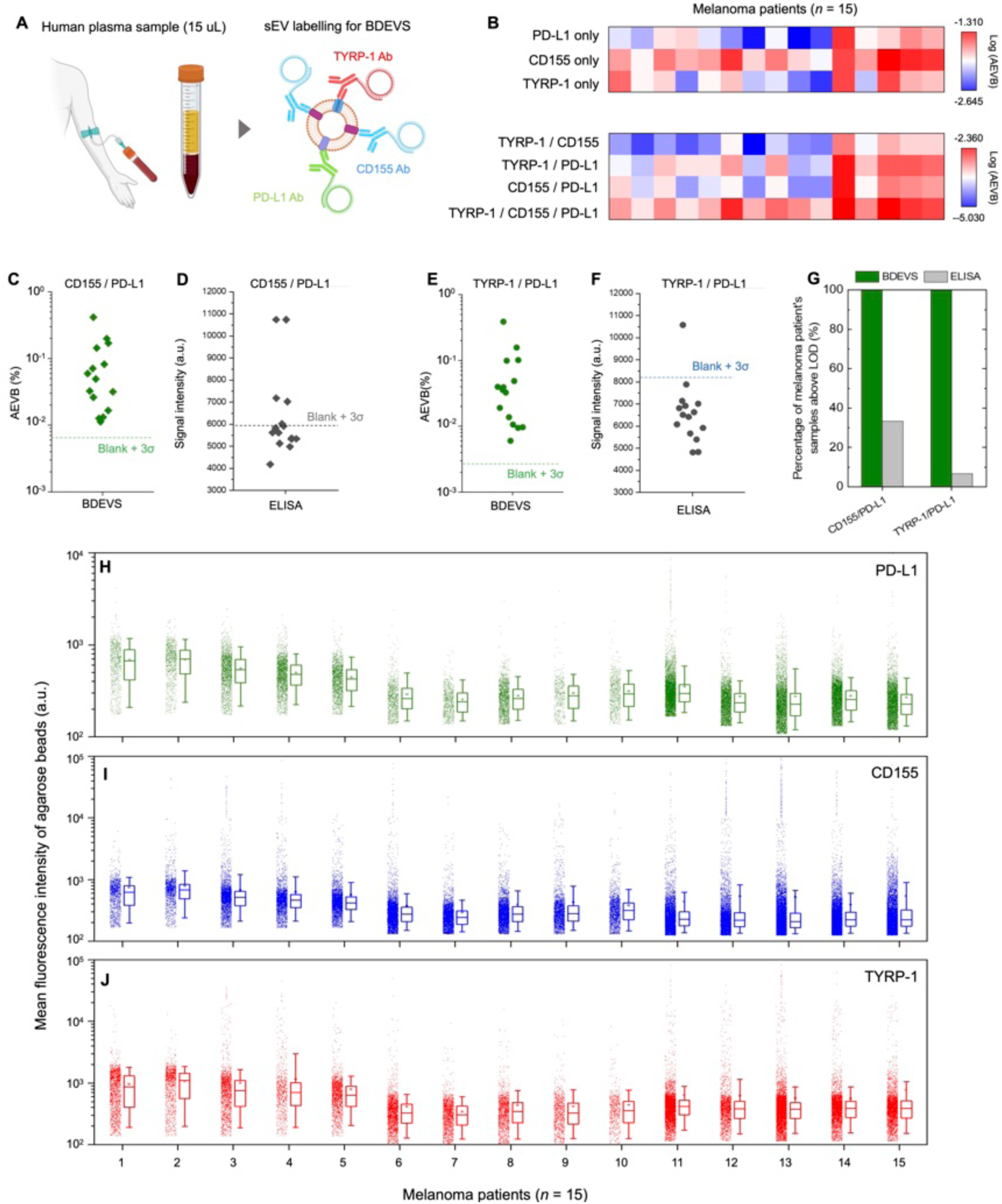
Analysis of single EVs in human plasma specimens from subjects with melanoma. (A) Schematic of workflow to analyze single EVs from human melanoma plasma samples for TYRP-1, CD155, and PD-L1 expression using BDEVS. (B) Heat map showing the AEVB of different populations of EVs from 15 melanoma patients’ plasma. Analytical performance to measure (C-D) CD155 / PD-L1 or (E-F) TYRP-1 / PD-L1 dual positive EVs from melanoma patients’ plasma were compared between BDEVS and ELISA. For ELISA, capture antibody targets CD155 or TYRP-1, and detection antibody is specific for PD-L1. (F) The ratio of melanoma patient’s plasma samples above LOD was compared between BDEVS and ELISA. (G) The ratio of patient’s plasma samples detectable by each method was plotted. (H-J) The quantitative expression of biomarkers per biomarker-positive EV was compared among melanoma patients.

## Discussion

In this study, we have developed a novel method to quantitatively analyze single EVs with multiplexing capabilities, high throughput, and ultra-sensitivity, both to single molecules on an individual EV and to rare EVs suspended in complex clinical specimens, using an agarose droplet-based RCA technique. Our approach utilizes cleavable chemistry to label RCA templates to single EVs, which are then digitally distributed over agarose beads for single EV / single-molecule level quantification. The RCA products retained within the agarose beads are analyzed by flow cytometry and sorted using FACS. Our results also highlight the versatility of the method for multiplex analysis by targeting multiple biomarkers, which can be further expanded through rational design of RCA templates. We demonstrated the potential of our platform for resolving EVs directly in plasma by quantifying immune markers PD-L1, CD155, and the melanoma tumor marker TYRP-1, on individual EVs directly in plasma from subjects with melanoma.

While a significant step forward, there are several opportunities to further develop our BDEVS technology. Currently, our platform can only quantify EVs positive for the surface markers targeted, which makes it impossible to directly quantify the total number of EVs. To address this current shortcoming in the technolgoy, the addition of lipid staining dyes or biotinylation of EVs followed by streptavidin-RCA template labeling could ensure accurate quantification of the total number of EVs input into our system. While high-throughput sorting can be achieved for downstream imaging, to perform digital quantification, the throughput is limited by imaging speed. We believe the incorporation of imaging flow cytometry can address this issue.[55,56] Additionally, because EVs are >30 nm, they are expected to be retained intact within our sorted agarose beads, making it possible for downstream analyses including RNA sequencing and mass spectrometry on subpopulations sorted based on multiparameter analysis of our BDEVS assay.

Increasing the multiplexing capacity of the assay could provide deeper insights into the interactions between different EV populations, such as those derived from macrophages and cancer cells, thereby advancing our understanding of cancer immunology and the tumor microenvironment. In this study, we analyzed ratio of immune marker positive EVs derived from melanoma cancer cells. It also has known that macrophage derived EVs play a significant role in cancer microenvironment.[59] With an increased multiplex capacity, analyzing the ratio of EVs derived from macrophage and cancer cell is expected to provide great advances for understanding cancer immunology. Furthermore, quantification of biomarker expression level of single EV derived from macrophage or cancer cell can provide great advances to understand EV’s role in cancer immunology.

## Methods

### Fabrication of the microfluidic droplet generator

SU-8 mold for a microfluidic droplet generator was designed and fabricated on a silicon wafer, and was silanized by vapor deposition of trichloro(1H,1H,2H,2H-perfluorooctyl)silane (Sigma Aldrich). PDMS was molded from SU-8 mold and bonded to glass slide after oxygen plasma treatment for 30 s. The channels were further silanized using 2% trichloro(1H,1H,2H,2H-perfluorooctyl)silane in HFE-7500 oil before use. Parallelized microfluidic droplet generators were fabricated using double side imprinting using a previously published method.[32,42] Briefly, an SU-8 replica mold “hard master” was generated with two layers, 50 μm height for the droplet generators and 200 μm height for outlet collection channels and vias were fabricated on a silicon wafer. An SU-8 mold for polydimethylsiloxane (PDMS) soft master was fabricated with 300 μm height on a silicon wafer. This mold was silanized using vapor deposition of trichloro(1H,1H,2H,2H-perfluorooctyl)silane for 2 h prior to being used to generate the soft master. From this mold, we created the soft master by curing 20:1 elastomer to crosslinker PDMS at 65 °C overnight. Using our hard master and soft master, we generated our microfluidic device by placing 8:1 PDMS sandwiched between the hard and soft master, and curing at 65 °C for 2 h. To facilitate the release of double sided patterned PDMS, both the hard master and soft master were silanized by vapor deposition of trichloro(1H,1H,2H,2H-perfluorooctyl)silane. The released double sided patterned PDMS device was washed with isopropyl alcohol, dried, and attached to a 3 mm thick 10:1 PDMS slab and then bonded to glass slide after 30 s of oxygen plasma treatment. Microfluidic channels were silanized with 2% trichloro(1H,1H,2H,2H-perfluorooctyl)silane for 30 min in HFE-7500 oil before use to render their surface hydrophobic.

### Preparation of DNA primer labelled antibody and circle DNA

Antibody was labelled with DNA primer using OYO link technology (AlphaThera) according to the provided protocol, which were bridged with S-S bond. Briefly, 1 µg of antibody was mixed with 1 µL of OYO linker – DNA primer (33 µM) and they were cross-linked under UV for 2 h. Labelling of DNA primer was validated by SDS-PAGE with 4-12% Bis-Tris gel (Invitrogen) (**Figure S11**). To prepare circle DNA, 5 ’phosphorylated DNA ogligomer was circularized using Circligase (Lucigen) under the manufacturer’s protocol. After that, non-circularized linear probes and their secondary structure were removed by Exonuclease I and III. Circularization was assessed by using 15% TBE-urea gel electrophoresis (**Figure S12**). Sequence information of DNA primer and circle DNA was summarized in **Table S2**.

### EVs labelling with RCA template

Mel-624-B7H1 cell line was cultured RPMI 1640 medium (Invitrogen) supplemented with 10% exosome-depleted FBS (Invitrogen). EVs from cell culture media were isolated using exosome isolation kit (Invitrogen) and resuspended in PBS. For plasma sample analysis, plasma samples were centrifuged 10,000g for 10 min and supernatant was diluted 10 times with 1% BSA. Both EVs from cell culture media and plasma samples were incubated with 5 nM of Ab-primer conjugates for 1 h at RT. EVs labelled with Ab-primer were purified using the qEV single column (70 nm, IZON) with 400 µL of elution volume. 10 µL of EVs labelled with Ab-primer was mixed with 10 µL of circle template (50 nM) were further incubated for 2 h at RT. EVs labelled with RCA templates were diluted to desirable ratio before use.

### Workflow of BDEVS

Ultra-low gelling temperature agarose (Sigma Aldrich) was dissolved in PBS to 1% (w/v) at 90 °C for 10 min. Molten agarose (0.5 ml/hr), 3x RCA mixture (1,200 U/mL phi29 polymerase, 0.75 mM dNTPs, 0.6 mg/mL recombinant albumin in 3x RCA reaction buffer) (0.5 mL/h), and EV-RCA template (0.5 ml/hr) was flown into the microfluidic device with BioRad oil for Evagreen (10 ml/hr) to generate 0.33% agarose droplet using a parallelized droplet generator. Agarose droplets were directly collected to a 1.5 mL tube in an ice bucket for the gelation. Then, 1.5 mL tube was incubated at 37 °C for 2 h for RCA reaction. Then, oil was removed after 300g centrifugation for 1 min. Remaining surfactant was further removed by using 10% PFO in HFE-7500. After removal of 10% PFO in HFE-7500 upon 300g centrifugation for 1 min, agarose beads were further washed with 1% Span80 (Sigma Aldrich) in hexane (Sigma Aldrich). Supernatant was removed after 300g centrifugation for 1 min, and agarose beads were washed three times with 0.05% PBST after 1,500g centrifugation for 2 min. RCA products in agarose beads were labelled with 10 nM of detection probes for 15 min at 37 °C. After washing three times with PBST, agar beads are analyzed using flow cytometry (Penn Flow Core) or fluorescence microscope (Leica). To concentrate agarose beads for imaging, they were sedimented in a PDMS well (diameter = 5 mm) for 30 minutes before imaging. For ELISA, 100 µL of 10x diluted plasma was analyzed and the assay result was compared with BDEVS.

## Author Contributions

The manuscript was written through contributions of all authors. All authors have given approval to the final version of the manuscript.

